# Structural and functional insights into QueC-family protein in QatABCD anti-phage system

**DOI:** 10.1101/2025.11.03.686418

**Authors:** Zirui Gao, Feixue Li, Hao Wang, Xi Liu, Ningning Li, Weijia Xiong, Weidong Ma, Demei Sun, Dingfei Yan, Qingqing Ma, Lei Xu, Yi Zhang

**Author notes:** These authors contributed equally (Zirui Gao, Feixue Li, Hao Wang). Correspondence (Y.Z.).

## Abstract

QatABCD is a widespread prokaryotic anti-phage defense system comprising four protein components, out of which QatC, a QueC-family protein is the signature component. QueC family proteins are nucleoside biosynthesis enzymes involved in the biosynthesis of queuosine, a 7-deazaguanine derivative. Recently, QueC-family proteins were shown to catalyze a deazaguanylation protein-nucleobase conjugation reaction in type IV CBASS antiphage defense. However, the mechanism of QatABCD, even the function of QatC in this system, remains unknown. Here, we demonstrate that QatBCD forms a complex in which QatD is highly flexible. Crystal structures of the QatBC complex in apo and ATP-bound form support a shared role for QueC-family proteins in targeting protein substrates for N-terminal modification as in type IV CBASS. We show that the QatB N-terminal loop, its binding with QatC and QatC catalytic site are essential for QatABCD defense *in vivo*, suggesting a modification might occur analogous to CBASS. These findings provide structural and functional insights into QueC-family protein in QatABCD system, suggesting the conserved mechanisms and critical roles of QueC-family in prokaryotic immunity.

## Introduction

7-Deazapurine and its derivatives, as nucleobase analogs, are important in critical cellular functions widely distributed in nature^1^, such as RNA and DNA modifications and secondary metabolites^2-5^. Among the 7-deazapurine derivatives, queuosine (Q) is one of the most structurally complex and evolutionarily conserved tRNA modifications^2,6^. Incorporated at the anticodon loop of tRNAs, Q plays crucial roles in optimizing translational efficiency and fidelity^1,2,6-9^.

While eukaryotes salvage its precursor Q from diet or microbiota^10^, prokaryotes can synthesize Q *de novo* through a multi-enzyme pathway^11-15^. QueC (7-cyano-7-deazaguanine synthase, renamed from ybaX) is essential in the early stages of the Q biosynthetic pathway, driving the ATP-dependent two-step transformation of CDG (7-caboxy-7-deazaguanine) to intermediate nucleobase preQ_0_^11,14,16,17^. As diverse QueC homologs increasingly identified in antiphage defense systems^18-20^, the canonical tRNA-modifying enzyme QueC is now recognized as a critical player in the virus-host arms race^21^. Recent studies on type IV cyclic oligonucleotide-based antiphage signaling systems (CBASS) revealed that the QueC homolog Cap9 was repurposed for nucleobase-protein conjugation in an immunity context. In these systems, Cap9 mediates a novel modification called NDG (N-terminal 7-amido-7-deazaguanine) on the cyclic dinucleotide synthase CdnD—a process essential for activating antiphage immunity^22^. Beyond the role in CBASS system, QueC domains are also found in various anti-phage defense operons, such as the QueC homolog QatC within the qatABCD system^19^.

The QatABCD system exhibits broad-spectrum phage defense capabilities. The system from *Escherichia coli* NCTC9009 has been identified to provide protection against phages P1, T3, and λ ^19^. Additionally, the system derived from *E. coli* strain 46-1 demonstrates low-level protection against T4(C), a mutant lacking gene 56 which encodes a cytosine methyltransferase^23^. Moreover, the qatABCD homologous system in *Pseudomonas aeruginosa* demonstrates antiphage activity against *Casadabanvirus* and *Pbunavirus* genera^24^. The qatABCD system in prokaryotes consists of four components: qatA is an ATPase, likely supplies energy, qatB has no recognizable known domain or motifs, qatC has a QueC domain and qatD is a Tat D family DNase^19,25^. However, the mechanism of QatABCD system remains unclear. Here, we determined the crystal structure of QatBC complex, combined with in vivo analysis and structure-guided biochemical analyses, we determine the structural and functional basis of the QueC-family protein QatC within the QatABCD anti-phage system. We show that QatC forms a stable complex with QatB, in which the N-terminal loop of QatB inserts into the catalytic center of QatC. Crystal structures of the QatBC complex in apo and ATP-bound states reveal that QatC retains a conserved QueC catalytic core with unique N- and C-terminal extensions that mediate specific interaction with QatB. Structure-guided mutational analyses further demonstrate that the catalytic residues of QatC, the zinc-binding motif, and the QatB N-terminal tail are indispensable for anti-phage activity. Together, our findings uncover that QatC likely acts as a functional analog of Cap9, mediating protein modification within the QatABCD system, and highlight the evolutionary diversification of QueC-like enzymes as a general strategy in prokaryotic immune defense.

## Results

### QatB interacts with both QatC and QatD

Previous study showed that all the four components are required for the anti-phage activity against λ phage when the QatABCD system from *E*.*coli* NCTC9009 (Fig. 1a) is expressed in *E. coli* BL21 cells^19^. To elucidate how QatABCD defends against phage infection, first we tested whether the four components interact with each other. To this end, we co-expressed the four components together, with His-tag on one out of the four proteins each time, respectively. Out of the four expression combinations, we found that QatB/C/D could form a complex, for which the best combination is His-tagged QatC but QatB and QatD without tags (Fig. 1b). The SEC-MALS (Size Exclusion Chromatography-Multi-Angle Light Scattering) analysis also indicated that QatB/C/D interacts with each other with a 1:1:1 ratio in the complex (Fig. 1c). Moreover, QatA does not interact with any of the other components. Next, we sought to analyze the interaction relationship of the three proteins in the QatBCD complex. The native gel results showed that complex can be formed either between QatB and QatC, or between QatB and QatD, but not between QatC and QatD (Fig. 1d). Furthermore, we confirmed this relationship through gel filtration assay (Extended Data Fig. 1a-c). This suggests that QatB might be a central component in binding to QatC and QatD. Taken together, protein complex can be formed between QatBC, QatBD and QatBCD in the system.

**Figure 1.**
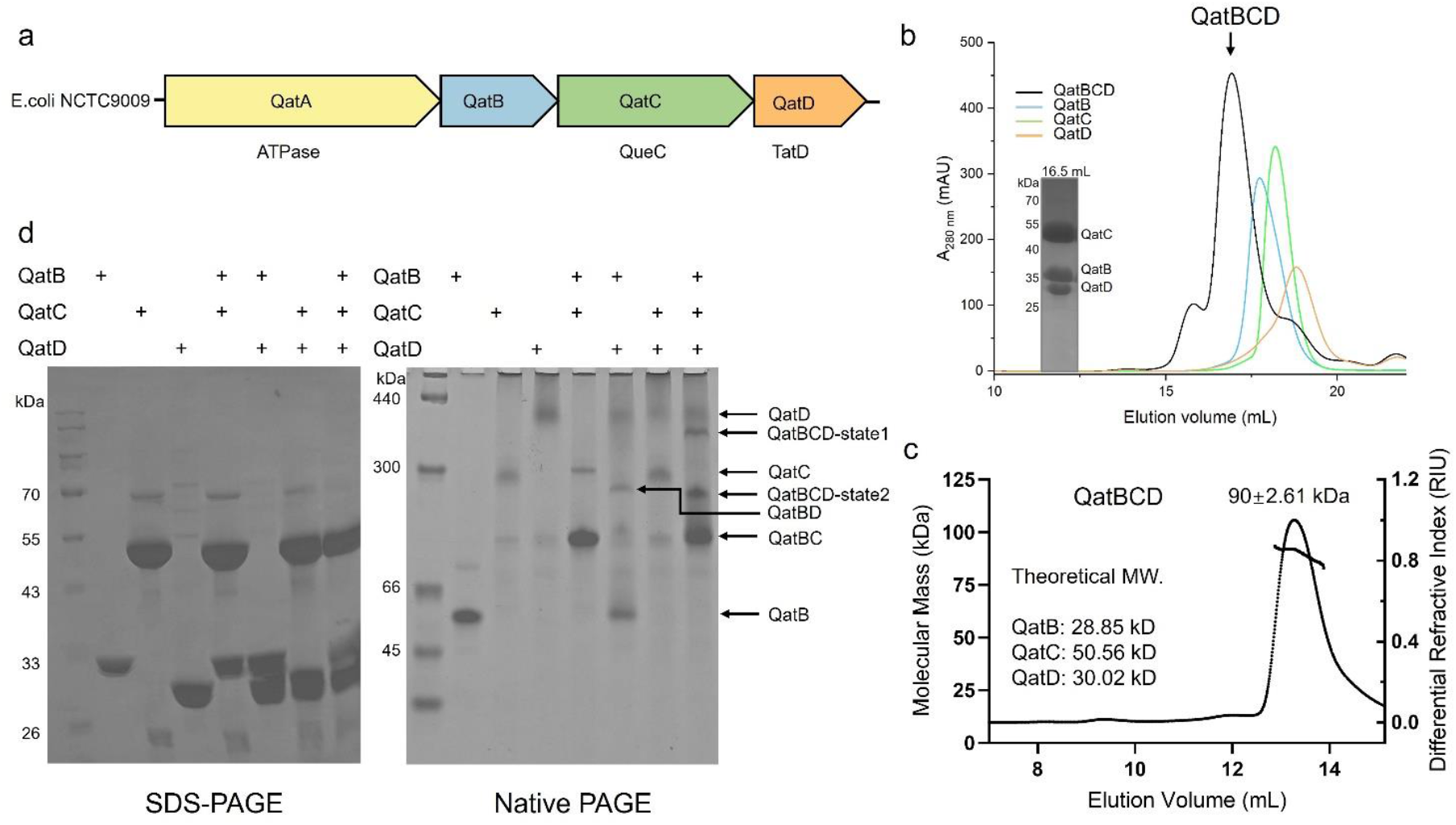
QatB, QatC and QatD form a stable complex. **a**, Schematic of the qatABCD system from *E*.*coli* NCTC9009. **b**, The gel filtration profiles of QatB, QatC, QatD and the mixture of them are shown. SDS-PAGE gel of the peak of the mixed sample is displayed. **c**, Static light scattering (SLS) studies of QatBCD complex. The calculated molecular weight of the main peak is shown above the peak. **d**, Native-PAGE showed the binding among QatB, QatC, and QatD. The results of the SDS-PAGE for the same samples are displayed on the left.

### Crystal structure of QatBC complex

To reveal the structural basis of QatBCD complex, we subjected the purified QatBCD complex for crystallization. However, despite extensive trials, there is only electron density for the QatBC complex in the crystal, but not QatD, suggesting that the QatD component might be flexible in this complex. Crystallization of the QatBD complex was also unsuccessful. Therefore, we finally solved the crystal structure of QatBC complex at 2.09 Å resolution for structural analysis (Extended Data Table 1). In the asymmetric unit, there are two 1:1 QatBC complex connected through the N-terminal β strand of QatC (Extended Data Fig. 2). However, PISA (Protein Interfaces, Surfaces and Assemblies) analysis^26^ indicated that QatBC is stable as a 1:1 complex, which is also supported by the SEC-MALS analysis of QatBC complex (Extended Data Fig. 1d). In the structure, QatB and QatC form a compact assembly with a large buried area of 4849.4 Å^2^ (Fig. 2a). In the complex, the QatC protein can be divided into three parts based on its QueC domain in the middle, that is, the C-terminal extension domain, the N-terminal extension domain and the QueC-like domain (Fig. 2b). Typically, the residues 148-370 of QatC constitute a core domain which is highly structurally similar to Q biosynthetic enzyme QueC from *Bacillus subtilis* (BsQueC) and Cap9 from *Rhizobiales* in type IV CBASS (Fig. 2c). Structural alignment of QatC with BsQueC and Cap9 also reveals overlapping active sites that are highly conserved across these enzymes^20^ (Extended Data Fig. 3a). The QueC-like domain includes a Rossman-like fold, consisting of five parallel β-strands with eight α-helices surrounded at both sides (Extended Data Fig. 4a). There is a conserved pyrophosphate-binding SXGXDS motif (S156-G158-D160-S161) located in the loop linking β5/α4 and at the top of α4 (Extended Data Fig. 3a). A highly conserved CxxCxxC motif (residues C352, C355, C358) and another cysteine C332 together coordinates a zinc atom (Fig. 2d). Mutation of the four cysteine residues also severely decreased anti-phage activity of the system (Fig. 2e), suggesting the possible important role of these residues in stabilizing the structure. In addition, the N-terminal extension of QatC comprises four parallel β-strands β1–β4 flanked by three α-helices α1–α3. The C-terminal extension of QatC is mainly composed of three α-helices α12–α14, which form interactions with QatB from one side (Fig. 2b and Extended Data Fig. 4a).

**Figure 2.**
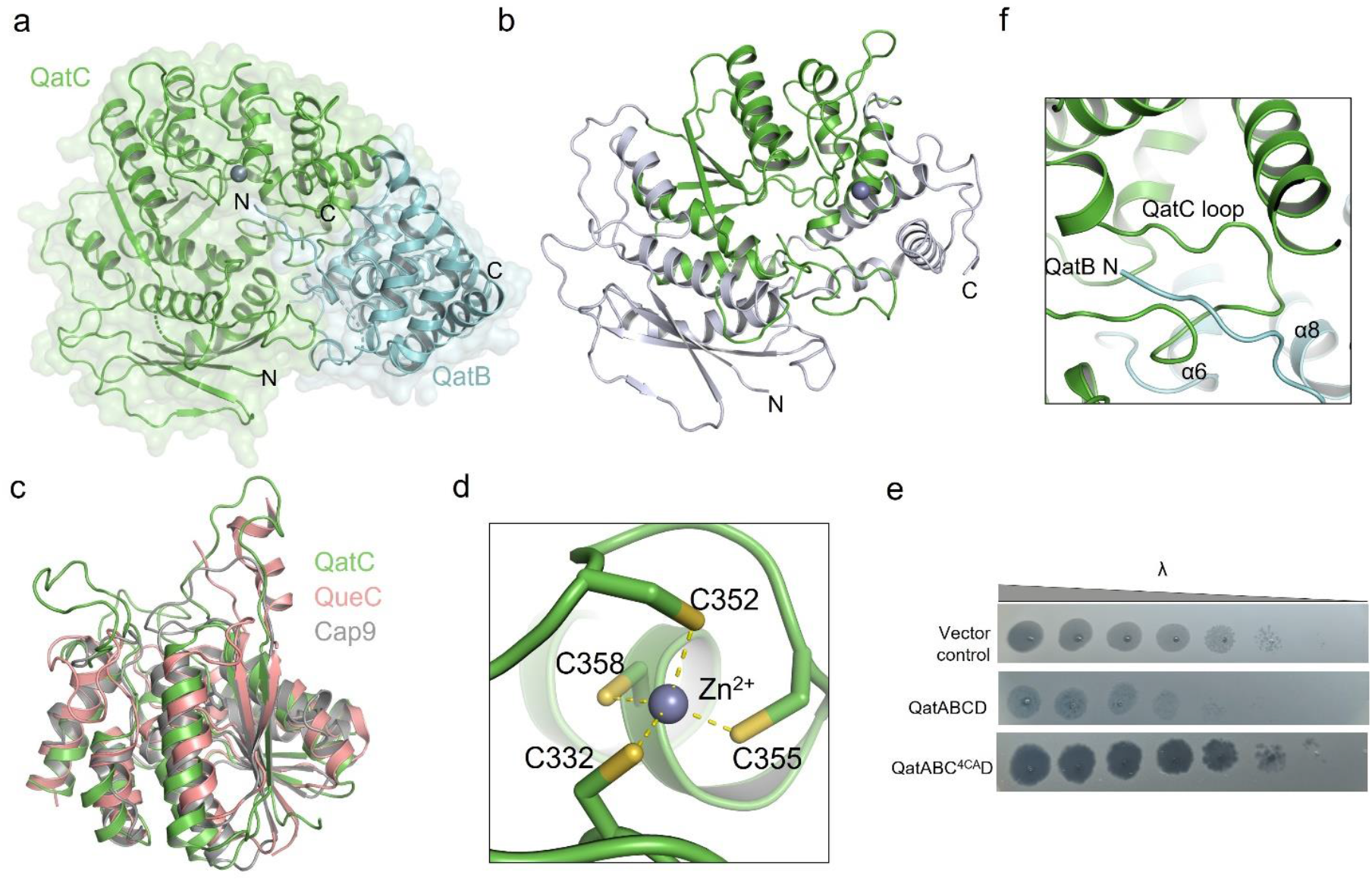
X-ray crystal structure of QatBC complex. **a**, The structure of QatBC complex is shown in the cartoon and surface model. **b**, Overall structure of QatC. The QueC-like domain is colored green. **c**, Structural superimposition among QatC, BsQueC (PDB: 3BL5) and Cap9 (PDB: 9NTO). Three proteins are displayed as green, pink, and gray, respectively. **d**, Detail diagram of zinc ion binding site of QatC. **e**, Validation of the zinc-coordinating residues of QatC by plaque assay. 4CA represents mutation of the four Cys residues shown in d to Ala. **f**, A close view of the binding of the QatC loop between α6 and α8.

QatB is a globular all α-helical structure (α1–α10) with a long N-terminal extension loop (Fig. 2a and Extended Data Fig. 3b and 4b). No similar entries with high Z scores returned from Dali search using QatB as a query^27^. All the three domains of QatC are involved in interaction with QatB. The most notable feature of the interaction is that the QatB N-terminal loop is inserted into the putative active site of QatC (Fig. 2a), suggesting that QatB might work as a substrate of QatC, like what the CD-NTase CdnD does in Cap9^22^. While both QatB and CdnD inserts its N-terminal loop into the QueC-like domain of QatC and Cap9 respectively, no structural homology exists between QatB and CdnD (Extended Data Fig. 5). In addition, the QatC residues N263–H280 form a long loop between QatC α6 and α8, which plugs into the surface cavity on QatB formed by its α6 and α8 (Fig. 2f). More detailed interactions between QatB and QatC will be discussed below. Together, the structure of QatBC complex displays how they interact with each other and suggests the possible role of QatB as a substrate of QatC.

### Catalytic site of QatC

Since QueC enzymes catalyze a two-step ATP-dependent reaction in which CDG (7-carboxy-7-deazaguanine) reacts with ATP as the first step, we moved on to incubate QatBC complex with both CDG and ATP, and solved a 2.66 Å X-ray crystal structure (Extended Data Table 1). However, there is no electron density for CDG but only ATP, and there is no variation in the density of the N-terminus of QatB either. This suggests that CDG might not have reacted with ATP in QatC, therefore we turned to analyze the binding pocket of ATP in QatC. Binding of ATP does not show significant impact on the conformation of the QatBC complex compared to the apo complex, with an RMSD of 0.308 Å among 656 Cα atoms (Fig. 3a). Investigation into the active site of QatC reveals conserved ATP binding and catalytic residues among QatC homologs as well as QueC and Cap9 (Fig. 3b and Extended Data Fig. 3a). Specifically, QatC F235 displays a marked conformational change compared with its apo state, to form π-π stacking with the adenine base of the ATP molecule. Moreover, the pyrophosphate-binding SXGXDS motif (S156-G158-D160-S161) together coordinates the β- and γ-phosphates of ATP in the structure (Fig. 3b). In addition, the ribose group of ATP is also coordinated by a hydrogen bond from carbonyl oxygen atoms of P255 (Fig. 3b). Finally, the α-phosphate group is also stabilized by electrostatic interactions from the N-terminal G2 of QatB.

**Figure 3.**
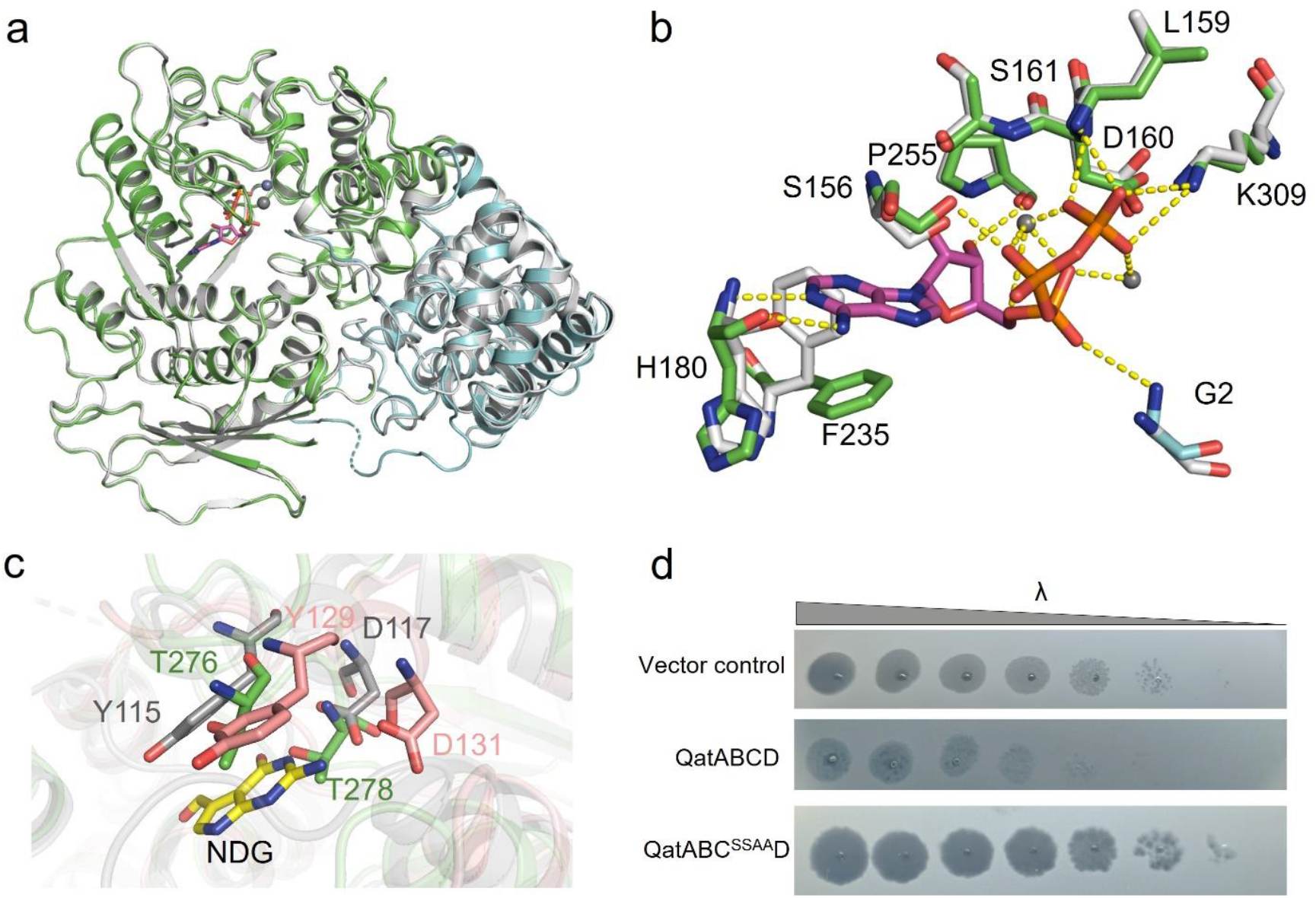
ATP-bound state structure of QatBC complex. **a**, Structural comparison between QatBC in its apo (gray) and ATP-bound form (green). **b**, Structural alignment of the binding pocket of ATP between QatBC in its apo (gray) and ATP-bound form (green). Hydrogen bonds are shown as yellow dashed lines. **c**, Structural comparison of the binding pocket of NDG among QatC (green), QueC (pink) and Cap9 (gray). **d**, Validation of the catalytic activity of QatC by plaque assay. SSAA represents QatC mutation S156A/S161A.

Next, we investigated the potential QatC substrate binding site, since there is no density for the CDG molecule. In the reported Cap9-CdnD structure, the N-terminus of CdnD is modified by the NDG nucleobase, while there is no density for the CDG molecule either^22^. Notably, the NDG modification is stabilized by the surrounding residues Y115 and D117, which are also conserved in BsQueC (Y129 and D131), suggesting the importance of these two residues in coordination of CDG and stabilization of the NDG moiety of the product. However in QatC structure, they are replaced by T276 and T278, respectively (Fig. 3c). Moreover, these two Thr residues are also conserved among QatC homologs (Extended Data Fig. 3a), suggesting their conserved function among QatC proteins, as well as the possibility of another molecule distinct from CDG as the substrate of QatC proteins. Mutations of QatC S156 and S161 significantly reduced anti-phage activity, indicating that the catalytic activity of QatC is indispensable for the function of the system (Fig. 3d). To investigate whether QatC is a functional QueC enzyme, we incubated EcQatC with ATP in the presence or absence of CDG and then analyzed the product through High Performance Liquid Chromatography (HPLC). Despite extensive trials, the results showed that EcQatC does not exhibit obvious ATP pyrophosphatase activity either in the presence or absence of CDG (Extended Data Fig. 6). Moreover, no obvious ATP pyrophosphatase activity was observed either when QatC was replaced by QatBC or QatBCD complex (Extended Data Fig. 6). Together these results indicate that QatC catalytic activity is essential for the immune function of the system but its substrate apart from QatB still awaits further studies.

### QatB N-terminal tail is essential for anti-phage activity

As mentioned above, the N-terminus of QatB is inserted into the active site of QatC. Like CdnD in the Cap9 active site, the electron density also starts from the second Gly residue of QatB, suggesting that the M1 residue was removed by post-translational proteolysis in bacteria, especially occurring when the second residue carries a small sidechain such as glycine^28^. Specifically, the N-terminal loop of QatB is stabilized by multiple hydrogen-bond and hydrophobic interactions from the active site cavity of QatC (Fig. 4a). The similar position of the N-terminal loop of QatB and that of CdnD in QatC and Cap9 (Fig. 4b), respectively, suggests that the G2 residue of QatB might also undergo modification by QatC. Notably, the N-terminal four residues “MGTS” of QatB are highly conserved among QatB homologs (Extended Data Fig. 3b), suggesting the importance of the N-terminus of QatB. Consistently, deletions of residues 14-20, mutation of G2 to aspartate or adding an N-terminal His-tag before QatB M1 all severely decreased the anti-phage activity of the Qat system (Fig. 4c). Despite extensive trials, we failed to detect NDG modification on QatB either using QatB proteins that are bacterially expressed or from *in vitro* reaction via mass spectrometry (Extended Data Fig. 7). Taken together, these results collectively define the essential role of QatB N-terminus in anti-phage activity.

**Figure 4.**
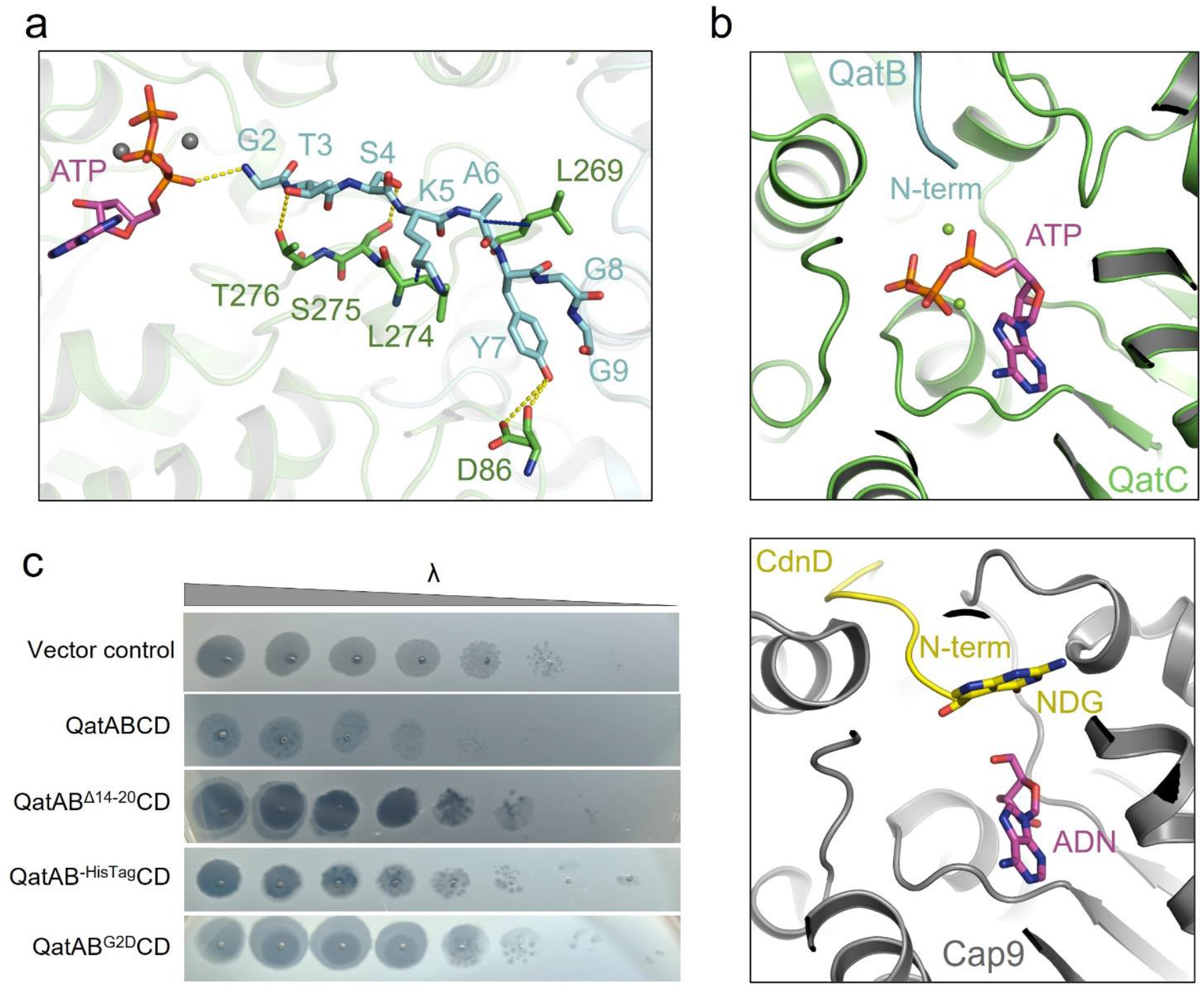
The N-terminus of qatB is essential for phage resistance. **a**, Detailed binding between the QatB N-terminus and QatC. Residues involved in binding are shown as sticks. Hydrogen bonds and hydrophobic interactions are shown as yellow and blue dashed lines. **b**, Structural comparison between QatB (cyan)-QatC (green) complex and CdnD (yellow)-Cap9 (gray) complex. **c**, Validation of the function of the N-terminus of QatB by plaque assay.

To investigate whether QatBC interaction is essential for anti-phage activity, we analyzed their interface in detail. Notably, a network of hydrophobic and electrostatic interactions is observed at their interface (Fig. 5a-e), in which hydrophobic residues of QatC (A404, P422, F282, L269, P268, and F415) engage in van der Waals interactions with hydrophobic residues of QatB (V218, I214, P147, A6, I208, M200, L138, and A165). Additionally, positively charged residues of QatC (R454 and R412) form salt bridges with negatively charged residues of QatB (D212, D166, and E162). To confirm the roles of these residues in QatBC interaction, we introduced point mutations on these critical interfacial residues and purified mutant QatBC complexes (Fig. 5f). Notably, the reduced intensity of the QatB band in SDS-PAGE gels (compared to wild-type complex) indicated disrupted QatBC complex formation. Consistently, anti-phage activity was also decreased by these QatB/C mutations (Fig. 5g), suggesting that these interfacial interactions are indispensable for the anti-phage activity of QatABCD system.

**Figure 5.**
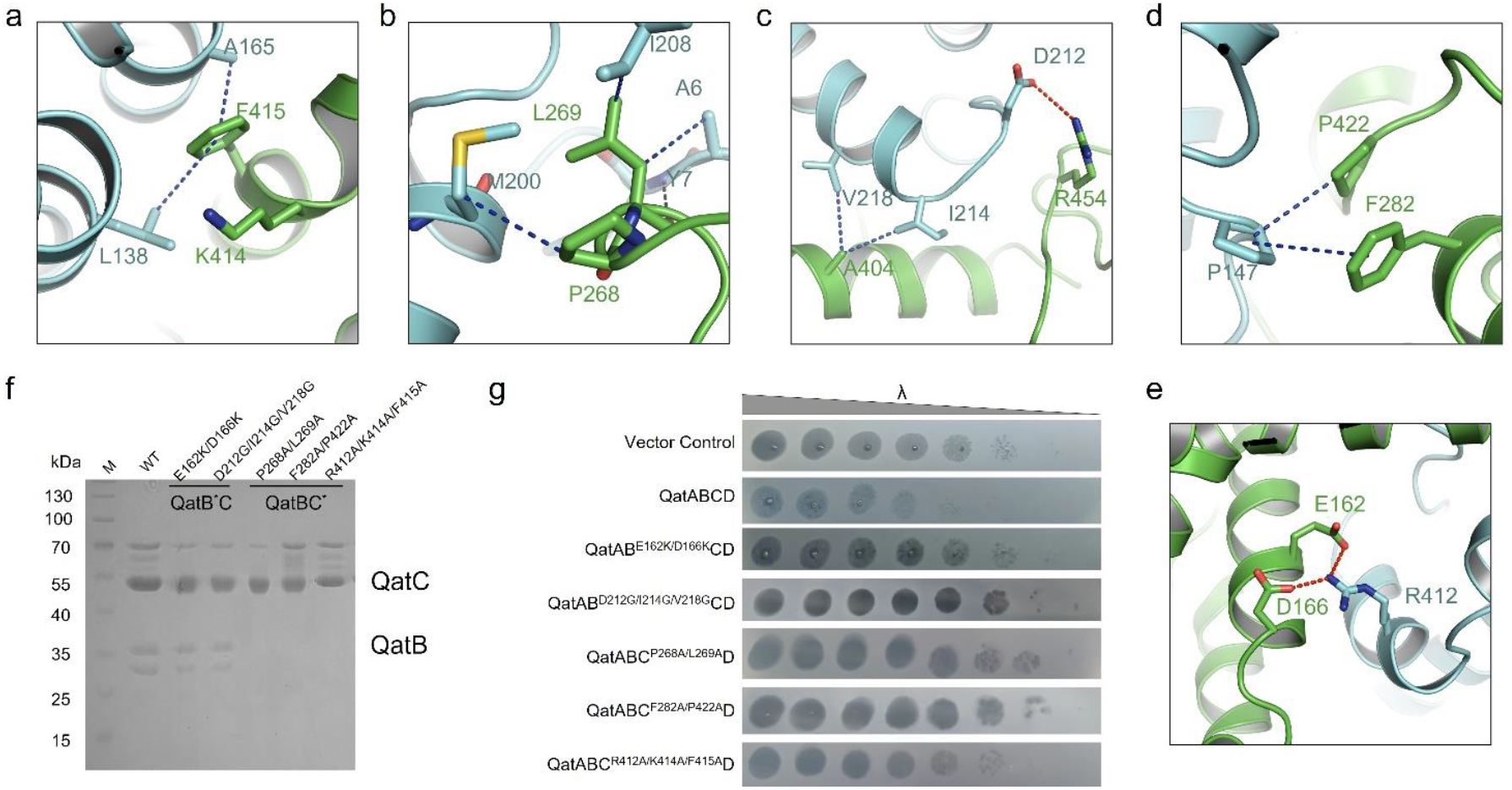
Interface interactions of QatBC are essential for activity of qatABCD system. **a-e**, Detailed interactions of QatBC complex are shown. Blue dashed lines represent hydrophobic and red dashed lines represent electrostatic interactions. **f**, Purified QatBC WT and mutant complexes were subjected to SDS-PAGE gel, showing the binding ratio between QatB and QatC, in which QatC was N-terminally His-tagged. **g**, Validation of the functions of the interface residues of QatBC by plaque assay.

### Insights into QatD in the QatBCD complex

Since QatD cannot be resolved through structural studies of the QatABCD complex, we used AlphaFold3 to predict a QatBCD complex (Extended Data Fig. 8a). In the predicted structure (interface predicted template modeling (ipTM) score of 0.61 and a predicted template modeling (pTM) score of 0.68), interestingly, QatC and the main body of QatB are in the same position as they are in the crystal structure, except that the QatB N-terminal flexible loop mainly interacts with QatD but not QatC, despite low confidence for this loop region (Extended Data Fig. 8a-b). Interestingly, in the structural alignment of the top five predicted structures of QatBCD, QatC and the main body of QatB overlaps well with their solved structure, but the position of QatD varies greatly (Extended Data Fig. 8c). This is consistent with our finding based on the crystal structure that QatD binding is highly flexible in the QatBCD complex. However, in each predicted structure, the QatB N-terminal flexible loop is involved in QatD binding (Extended Data Fig. 8b). To investigate the function of this loop in QatD binding, we generated a QatB Δ1–46 truncated mutant and native gel assay showed that this truncated mutant lost QatD binding (Extended Data Fig. 8d). Interestingly, this QatB truncated mutant still retains QatC binding (Extended Data Fig. 8d), suggesting that the other interfaces other than those involving QatB N-terminal loop play a key role in QatBC binding. The function of QatD in this system and its binding by QatB N-terminus still awaits further study. Together, these results suggest that QatD is flexibly bound in the QatBCD by the N-terminus of QatB.

## Discussion

In this study, we mainly characterize the QatBC complex in the QatABCD system through structural biology, biochemistry and *in vivo* studies. As the signature component of this system, QatC is a QueC-domain protein, whose catalytic activity is essential for the anti-phage activity of the system. Together with the recently published study into type IV CBASS^22^, this work illustrates the role of QueC-family proteins as key catalytic components in prokaryotic defense systems. Moreover, these studies also extend the substrate scope of QueC-family proteins beyond nucleic acids^29^.

Crystal structures of QatBC complex reveal a series of essential elements in the anti-phage activity of QatABCD, including the catalytic active site of QatC, the Zinc-coordinating residues of QatC, the N-terminal flexible loop of QatB and the interface between QatB and QatC, which have been proved by phage plaque assays^30^. Interestingly, the structure of QatBC was obtained coincidentally when we crystallized the QatBCD complex. Predicted structures of QatBCD support the notion that QatD is flexibly bound in the complex and suggest that QatB N-terminus is responsible for this binding, which was further confirmed by native gel assays. Previous study indicated that mutation of the putative active site of QatD decreases the anti-phage activity of QatABCD^22,23^. However, the exact role of QatD in the system needs further study. During the preparation of our manuscript, the structure of QatBC has also been reported by two other studies^31,32^. Interestingly, one of the studies reported that QatA and QatD in the *Pseudomonas aeruginosa* qatABCD system are not required for the anti-phage activity^31^. However, these two genes in the *E. coli* 46 NCTC9009 system have been confirmed to be required for anti-phage^19^. This suggests that for systems from different species distinct factors are needed for anti-phage activity.

It remains unknown what exact reaction is catalyzed by QatC. The crystal structure of QatBC suggests that QatB meets the conditions to be a substrate of QatC: its N-terminus inserted into the catalytic center of QatC and in the same context as CdnD in the Cap9 catalytic center. However, despite extensive trials, we could not reconstitute the activity of QatC using CDG, ATP and QatB. It suggests that QatC either needs a phage trigger to be activated or utilizes a substrate distinct from CDG. Future studies into this will help characterize the nature of the modification of QatB N-terminus and its role in anti-phage activity. While the complete mechanism of qatABCD-mediated defense remains unsolved, our results illustrate that qatABCD utilizes a mechanism in which QueC-family also mediates protein-ligand conjugation.

## Methods

### Bacterial strains and phages

The E. coli BL21 (DE3) strains were grown in Lysogeny broth (LB) medium at 37°C both with aeration at 225 r.p.m. When indicated, ampicillin was used to maintain the pBAD plasmid. Gene expression was induced by the addition of 0.2% L-arabinose.

The E. coli BL21 (DE3) strain was used for recombinant protein overexpression and grown in Lysogeny broth (LB) medium. The cells were grown at 37 °C until OD_600nm_ reached 0.8 and then induced at 18 °C for 12 h.

### Protein expression and purification

The *qatABCD* gene was bought from AddGene. The full-length *qatA/qatB/qatC/qatD* gene was amplified by PCR and cloned into a modified pET28a vector in which the expressed protein contains a His6 tag or His6-SUMO tag. The full-length *qatBC* gene was amplified by PCR and cloned into a modified pRSFDuet vector in which the expressed QatC protein contains a His6 tag. The *qatA* and *qatD* were respectively cloned into the two multiple cloning sites of a pETDuet vector without tag. The mutants were generated by two-step PCR and were subcloned, overexpressed and purified in the same way as for the WT protein. All of the proteins were expressed in E. coli strain BL21 (DE3) and induced by 0.2 mM IPTG when the cell density reached an OD600 of 0.8. After growth at 18 °C for 12 h, the cells were collected, resuspended in lysis buffer (50 mM Tris-HCl pH 8.0, 300 mM NaCl, 10 mM imidazole and 1 mM PMSF) and lysed by sonication. The cell lysate was centrifuged at 20,000g for 50 min at 4 °C to remove cell debris. The supernatant was applied onto a self-packaged Ni-affinity column (2 ml Ni-NTA, Genscript) and contaminant proteins were removed with wash buffer (50 mM Tris pH 8.0, 300 mM NaCl, 30 mM imidazole). The fusion protein were elution with elution buffer (50 mM Tris pH 8.0, 300 mM NaCl, 30 mM imidazole). The protein of QatA/QatB/QatC/QatD with His6–SUMO tag was digested with Ulp1 on the Ni-NTA column at 18 °C for 2 h after removing contaminant proteins with wash buffer. The protein was then eluted with wash buffer. The eluant of protein was concentrated and further purified using a Superdex-200 increase 10/300 GL (GE Healthcare) column equilibrated with a buffer containing 10 mM Tris-HCl pH 8.0, 200 mM NaCl and 5 mM DTT. The purified proteins were analysed by SDS–PAGE. The fractions containing the target protein were pooled and concentrated.

### Gel filtration assay

The QatB, QatC, QatD and QatBCD complex purified as described above were subjected to gel filtration analysis (Superdex-200 increase 10/300 GL, GE Healthcare). The QatB, QatC and QatD was incubated at a molar ratio of 1:1:1 overnight on ice before the gel filtration analysis in buffer containing 10 mM Tris-HCl pH 8.0, 200 mM NaCl, and 5 mM DTT. The assays were performed with a flow rate of 0.5 mL/min and an injection volume of 1 mL for each run. Samples from relevant fractions were subjected to SDS-PAGE and visualized by Coomassie blue staining.

### Crystallization

After molecular exclusion chromatography purification, the fractions containing the target protein were pooled and concentrated to 15 mg/mL. Both crystal forms of Protein were obtained using the sitting-drop vapor diffusion method at 18°C by mixing 0.8 μL of protein solution with 0.8 μL of reservoir solution. For the apo form, the reservoir solution contained 2 M ammonium sulfate. For the ATP-bound form, the reservoir solution was composed of 0.1 M ammonium sulfate, 0.3 M sodium formate, 0.1 M Tris pH7.8, 3% w/v γ-PGA (Na^+^ form, LM), 5% w/v PEG 4000. The protein was pre-incubated with 1 mM ATP and 2 mM MgCl_2_ on ice for 1 hour prior to crystallization. Before harvesting, the crystals were cryoprotected in the reservoir solution supplemented with 20% (v/v) glycerol and then flash-frozen in liquid nitrogen.

### Data collection, structure determination and refinement

All diffraction data were collected at the SSRF beamlines BL02U1 and were integrated and scaled using the HKL2000 package^33^. The initial model of Vs-3 was obtained using AlphaFold2^34^. All structures were further refined in PHENIX^35^ with non-crystallographic symmetry (NCS) and stereochemistry restraints. The final models were obtained after several rounds of refinement. All structural illustrations were generated using PyMOL (https://pymol.org/). Detailed statistics of data collection and structure refinement are summarized in Extended Data Table 1.

### DNA-digestion assays

The QatA/QatD/QatBCD (1 μM) and ssDNA/dsDNA/E.coli genome/λ genome (1.2 μM) were incubated at 37 °C for 30 min in 20 mM Tris pH 8.0, 150 mM NaCl and 5% glycerol. 1 mM ATP and 2 mM MgCl_2_ were added in the marked samples concurrently. The mixture was separated using 1% agarose gel and visualized by Coomassie blue R250 staining.

### SEC-MALS (Size Exclusion Chromatography-Multi-Angle Light Scattering)

Static light scattering experiments of QatA/QatA+ATP/QatBC/QatBCD were performed in 10 mM Tris-HCl pH 8.0, 200 mM NaCl, 5 mM DTT with a GE Healthcare Superdex-200 increase 10/300 GL size-exclusion column connected to the Wyatt DAWN HELEOS Laser photometer and Wyatt Optilab T-rEX differential refractometer^36^. Wyatt ASTRA 7.3.2 software was used for the data analysis.

### Native-PAGE assay

For the assessment of native-PAGE for the binding between proteins, each protein was pre-incubated at 20 μM at 18 °C for 20 minutes. Products of the reaction were analysed using 8 % native polyacrylamide gels and visualized by Coomassie blue staining.

### QatC activity assay

For testing the activity of QatC and its mutants, depyrophosphate reactions were performed in a 100 μL reaction volume containing 50 mM Tris pH 8.2, 50 mM KCl, 2 mM MgCl_2_, 2 mM CDG and 2 mM ATP if needed. After incubation at 37 °C for 60 min, the reactions were terminated by adding acetonitrile with a final concentration of 20 %. The products were filtered with a 0.22 μM filter and were subsequently used for HPLC experiments. The HPLC analysis was used an 100 Å AQ C18 column (4.6 x 250 mm; 5 μm; Bonnasil-BS). Chromatographic separation was performed at a flow rate of 1 mL/min under isocratic elution with 97 % 20 mM potassium phosphate buffer (pH 6.0) and 3 % acetonitrile. Detection was carried out at 254 nm.

### Ni-column pull down assay

20 μM each protein were first incubated together for 30 min at 37 °C, then the mixtures were incubated with Ni-NTA beads for 30 min at 4 °C. Buffer containing 300 mM NaCl, 50 mM Tris-HCl pH 8.0 and 30 mM imidazole was used to wash the beads. Samples of input and pulldown were separated using SDS–PAGE after washing three times.

### Phage plaque assay

For the plaque assay, the qatABCD operon cloned into the pBAD24 vector with any indicated mutations or deletions and transformed into Escherichia coli strain BL21(DE3). Cultures were incubated while shaking at 37 °C over night in LB medium. Bacterial culture was mixed with 0.8 % LB-agar to an OD600 of 0.06 supplemented 0.2 % L-arabinose, poured onto the surface of a 2.5 % LB-agar plate, and allowed to solidify at room temperature for 1 hour. 5 mL of top agar was used for 850×15mm round plates and 10 mL was used for 100×100mm square plates. Ten-fold serial dilutions of the phages were prepared, and 1.2 µL of each dilution was spotted onto the bacterial layer. After the spots had dried, the plates were inverted and incubated at 37 °C for 3 to 6 h before imaging.

### Mass spectrometric data collection and spectral deconvolution of intact proteins

QatBCD complex was expressed and purified using a N-terminal 6×His QatC tag as described above. Normalized protein samples were diluted to 1 μM in CH3CN/H2O (1:1) and analyzed by intact protein LC/MS using a Waters SYNAPT G2-Si Q-TOF system equipped with an ACQUITY UPLC HSS T3 VanGuard column. The mobile phase was a linear gradient of 5-99% ACN/water +0.1% formic acid. Peak integration and spectral deconvolution was performed using UniDec software.

### Mass spectrometric data collection and analysis of QatB N-terminal peptide

To determine the N-terminal modification of QatB, 5 μM QatB, 5 μM QatC, 1 mM ATP and 1 mM CDG were reacted in a buffer containing 25 mM HEPES pH 7.5, 100 mM KCl, 5 mM MgCl_2_ at 37°C for 1 hour. The complex was subjected to SDS-PAGE and the band corresponding to QatB was cut from the gel, resuspended in PBS and digested with chymotrypsin at 40:1 protein-to-protease ratio for 16 h at 37°C. Peptides were injected onto a UHPLC 3000 system coupled to a Thermo Scientific Orbitrap Exploris 480 mass spectrometer using a C‐18 analytical column (300 Å, 5 μm, Thermo Fisher Scientific, USA). The peptides were eluted with a gradient elution program at a flow rate of 0.300 μL/min. Mobile phase A consisted of 0.1% formic acid, and mobile phase B consisted of 100% acetonitrile and 0.1% formic acid. The mass spectrometer was operated in the data‐dependent acquisition (DDA) mode using Xcalibur 4.5.445.18 software. MS1 spectra were acquired at a mass range of 300–1800 m/z with a resolution of 60,000. The spray voltage was set at 2100 V, and the automatic gain control (AGC) target was set to 3e6. For MS2 scans, the top 40 most intense precursor ions were fragmented in the HCD collision cell at a normalized collision energy of 32% using a 0.4 Da isolation window. The dynamic exclusion duration was set to 15 s, and the AGC target was set to 1e5 while the maximum injection time was set to 100 ms^29^.

### Statistics and Reproducibility

For Figures 1b-d, 2e, 3d, 4c, 5f-g; Extended Data Figures 1, 6, 7, 8d, each experiment was repeated independently three times with similar results.

## Data Availability Statement

All data are available in the manuscript or the supplementary information. The crystal structure of QatBC and ATP-bound QatBC has been deposited in the Protein Data Bank under accession code 9WVJ and 9X6C, respectively.

## Author Contributions

Y.Z. conceived and supervised the project and designed experiments. Z.G., F.L., N.L., W.X., W.M., D.S. and L.X. purified the proteins and performed *in vitro* activity analysis and in *vivo* assays. H.W. and F.L. collected the diffraction data and solved the structures. Y.Z. wrote the original manuscript with the help of all the other authors.

## Acknowledgements

We thank the staff at beamlines BL02U1 and BL19U1 of the Shanghai Synchrotron Radiation Facility for their assistance with data collection; the staff at the Tsinghua University Branch of China National Center for Protein Sciences Beijing and S. Fan for providing facility support for X-ray diffraction of the crystal samples. Y.Z. is supported by National key research and development program of China (2024YFA0916903 and 2022YFC2104800) and the National Natural Science Foundation of China (32371329).

