## Supplemental Table 1 for "Structural and functional insights into QueC-family protein in QatABCD anti-phage system"

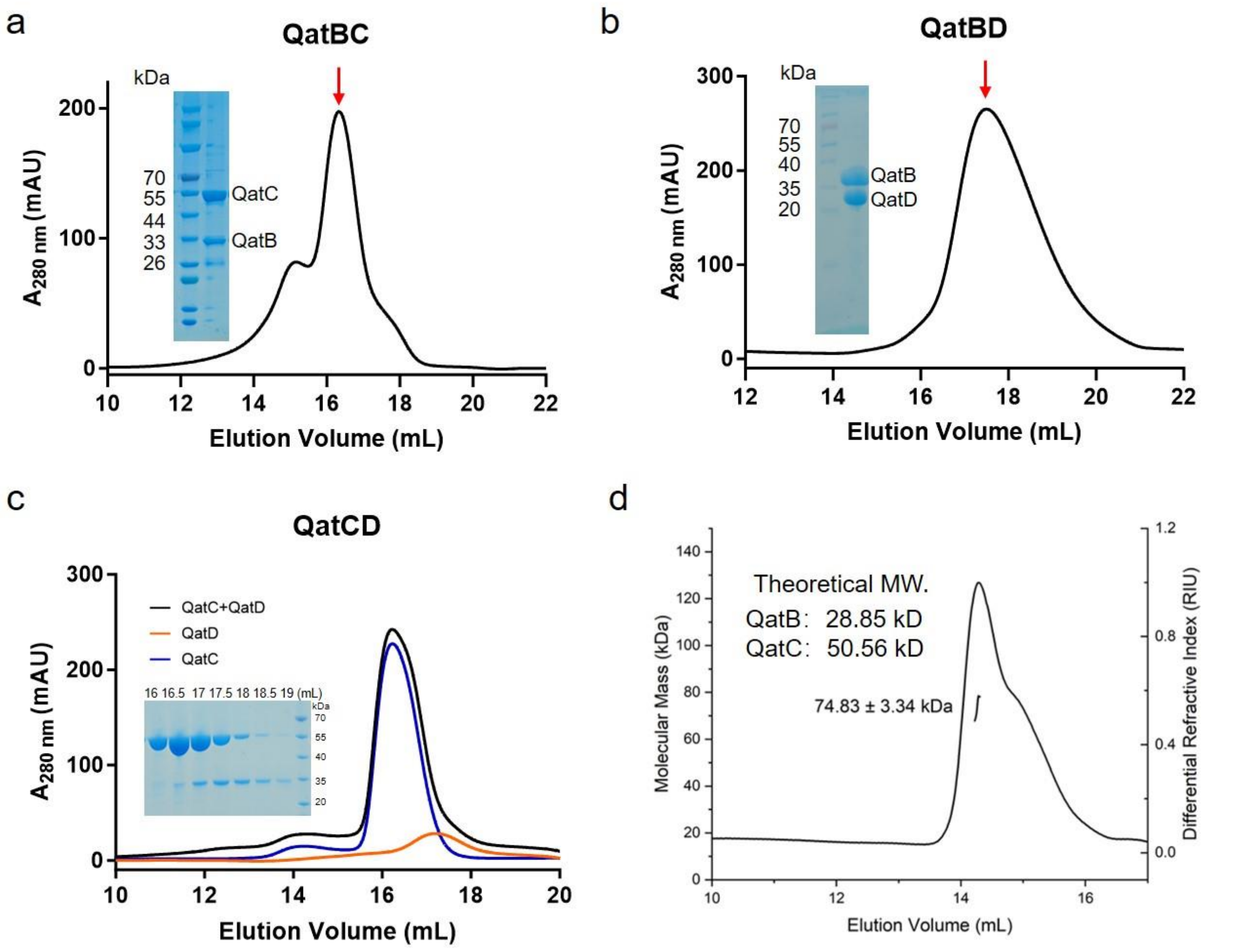


**Extended Data Fig. 1 QatBC and QatBD form a stable complex**

**a-b,** SEC assay of QatBC (a), QatBD (b). SDS-PAGE of peak tip sample is displayed.

**c,** The gel filtration profiles of QatC, QatD and the mixture of them are shown. SDS-PAGE gel of each sample tubes of QatC-QatD mixture is displayed.

**d,** Static light scattering (SLS) studies of QatBC. The calculated molecular weight of the main peak is shown beside the peak.

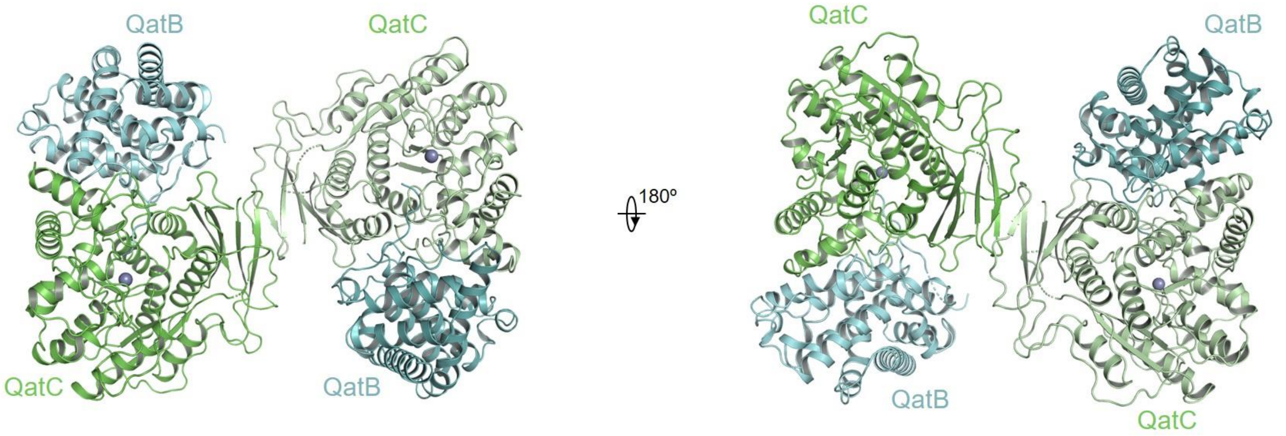


**Extended Data Fig. 2 Overall structure of QatBC complex.**
Overall structure of QatBC complex in the asymmetric unit. Two views are shown.

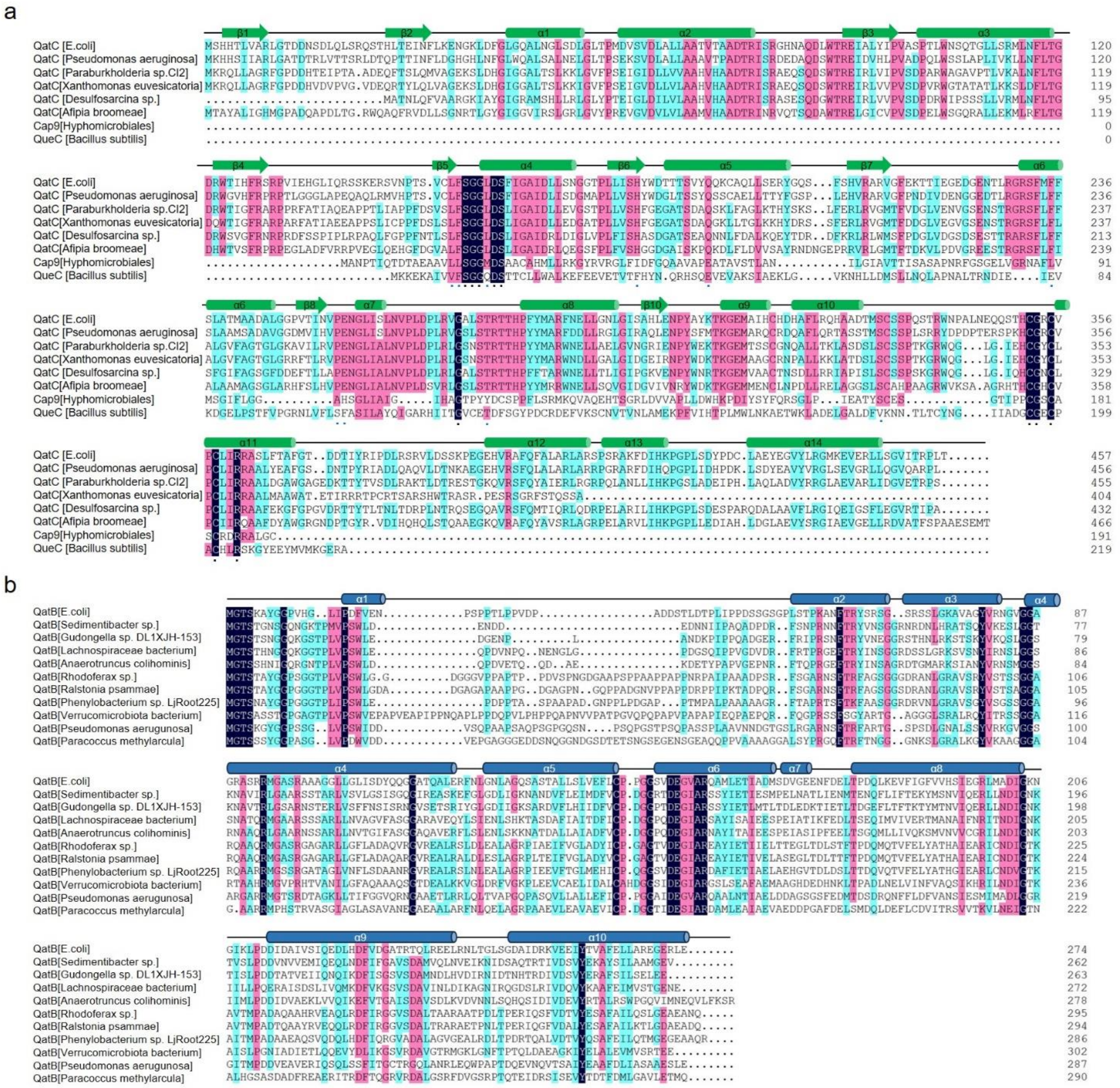


**Extended Data Fig. 3 Sequence alignment of QatB and QatC.**

**a,** Sequence alignment of representative QatC sequences, Rhizobiales sp. Cap9, and B. subtilis QueC. Secondary structure is annotated from *E.coli* NCTC9009 QatC structure in complex with QatB.

**b,** Sequence alignment of representative QatB sequences. Secondary structure is annotated from *E.coli* NCTC9009 QatB structure in complex with QatC.

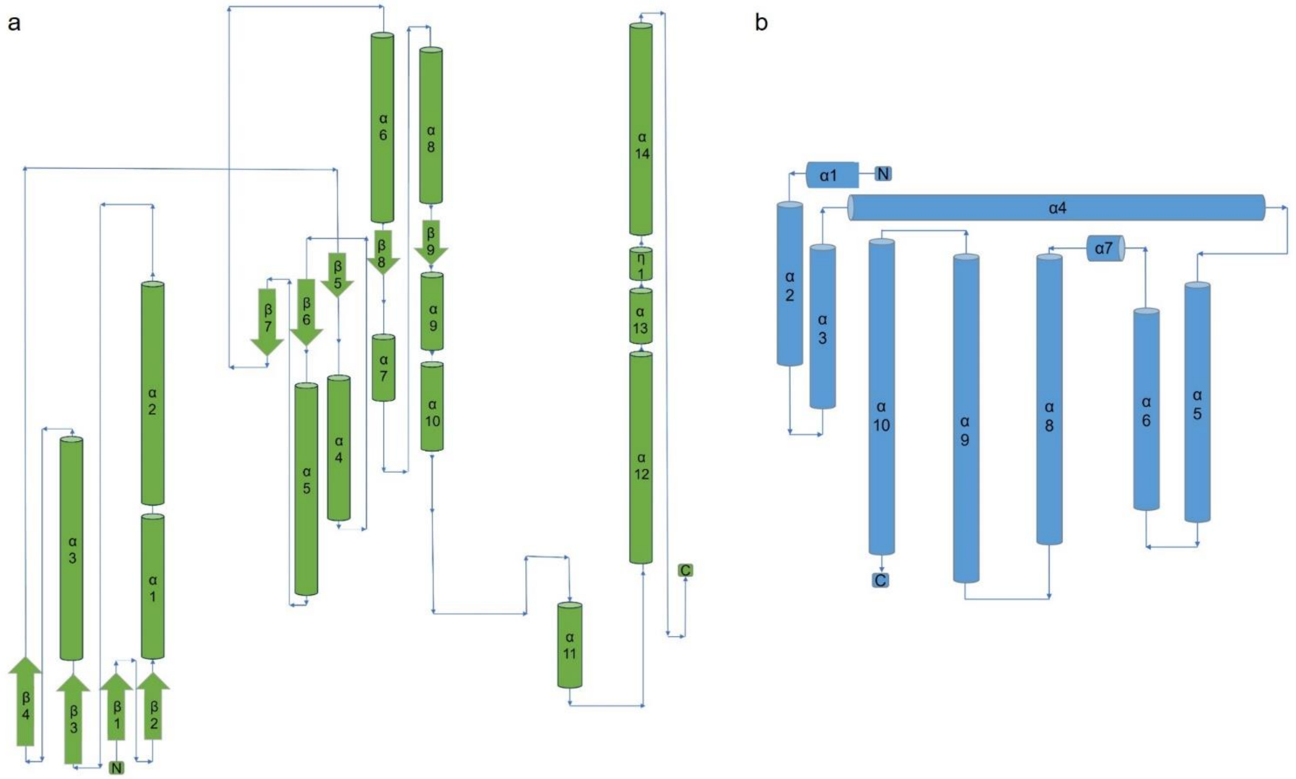


**Extended Data Fig. 4 Topology diagram of QatB and QatC.**

**a-b,** The diagram shows the secondary structure arrangement of QatC (a) and QatB (b). α-helices are represented as cylinders and β-strands as arrows, with their sequential order indicated. The topology is based on the crystal structure.

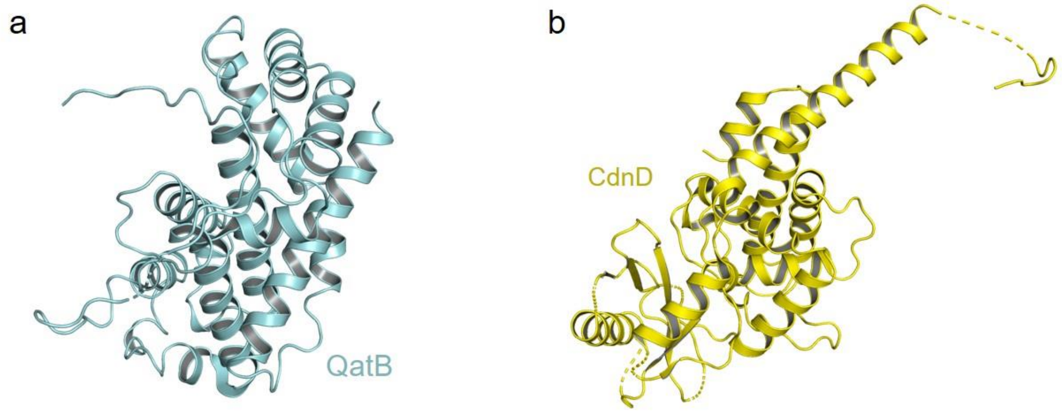


**Extended Data Fig. 5 There is no structural homology between QatB and CdnD.**
**a,** The structure of QatB in QatBC complex.
**b,** The structure of CdnD in Cap9-CdnD complex.

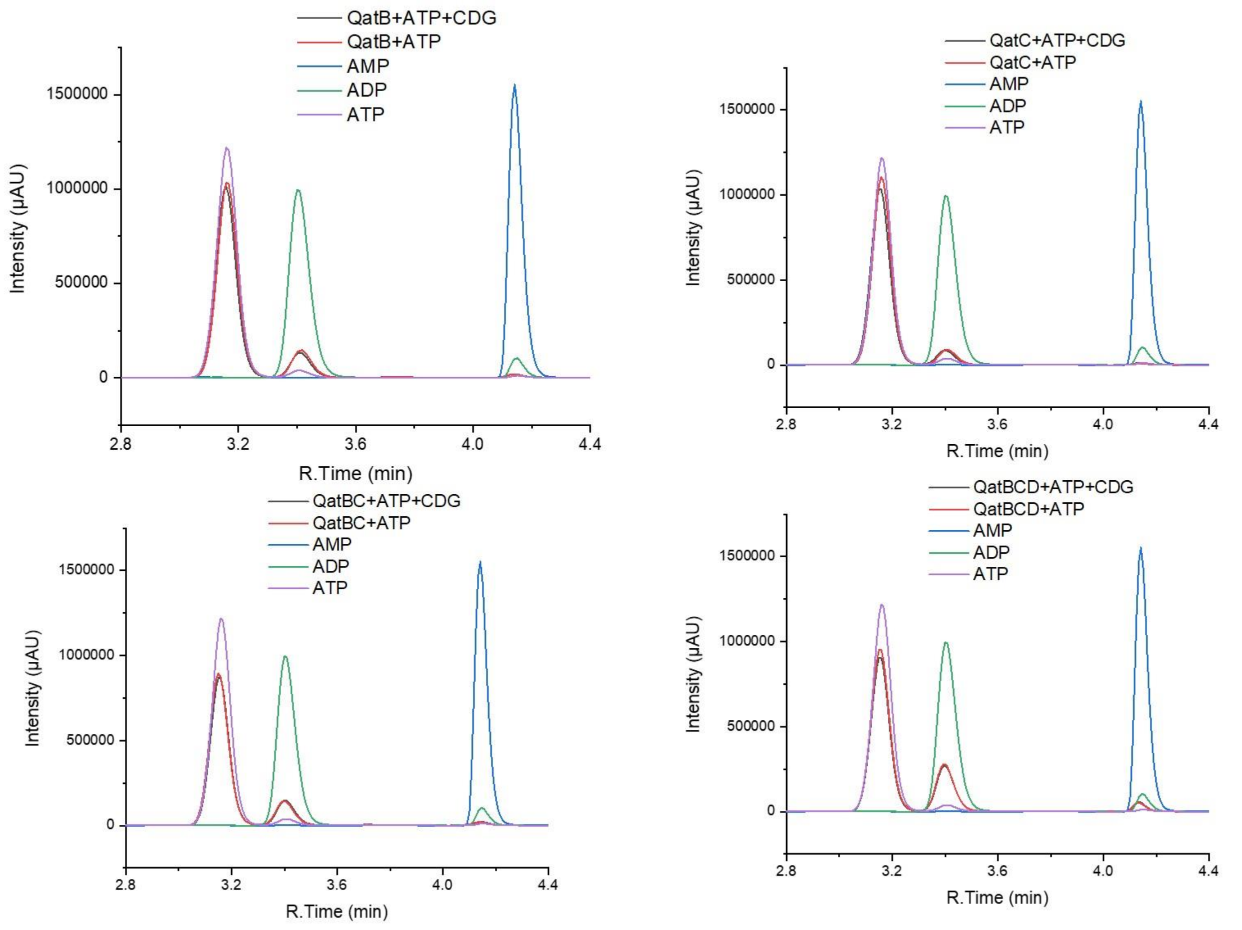


**Extended Data Fig. 6 HPLC analysis of enzymatic activity of QatC.**
HPLC analysis of enzymatic reactions with ATP and CDG in the absence or presence of QatB, QatC, QatBC and QatBCD. ATP, ADP and AMP standard is shown.

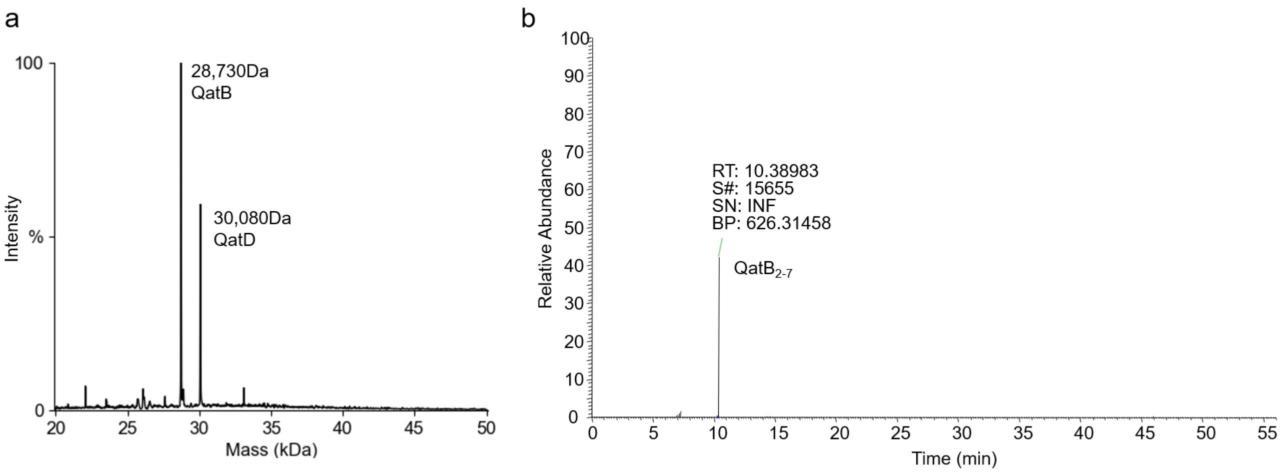


**Extended Data Fig. 7 LC-MS and chromatogram analysis of QatB**

**a,** Intensity of ion masses determined by intact LC-MS analysis and spectral deconvolution of the QatBCD complex.

**b,** Chromatogram of N-terminal peptides (residues 2 to 7: GTSKAY) from chymotrypsin-digested QatB. Mass-to-charge ratio (m/z) of peptide ions with charge 1+ is reported.

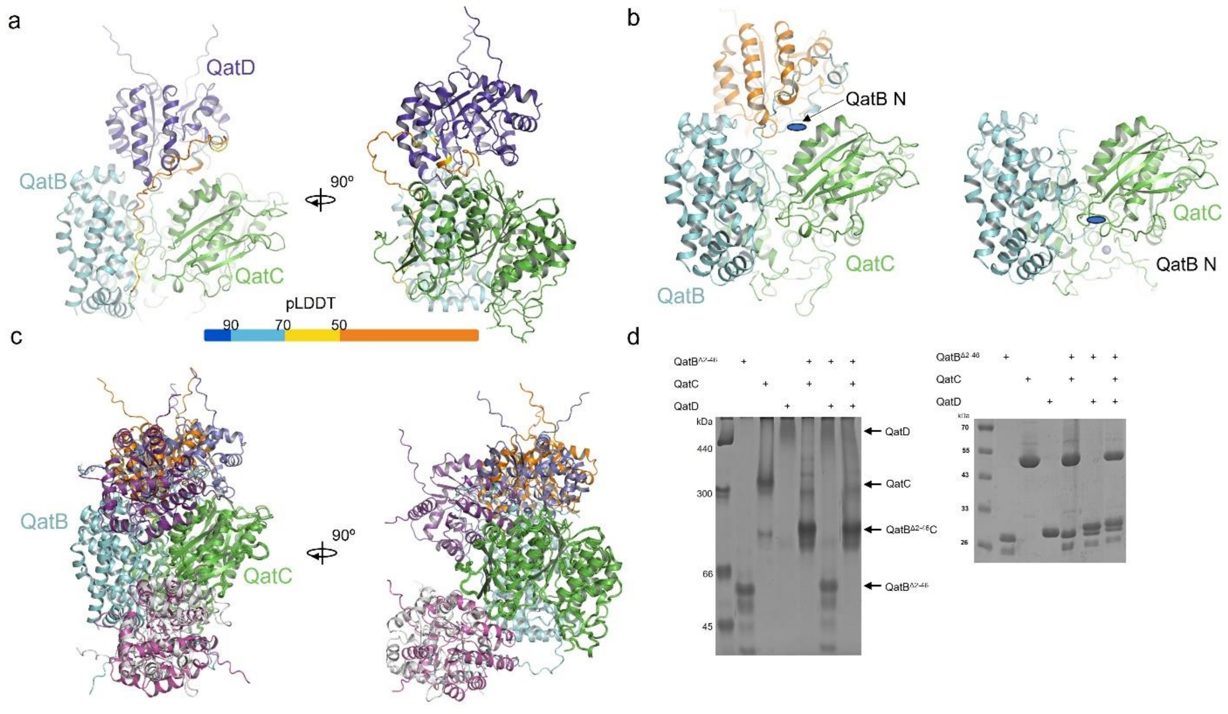


**Extended Data Fig. 8 The structure of QatBCD complex predicted by AlphaFold3**

**a,** The structure of QatBCD complex predicted by AlphaFold3. The N-terminal 51 residues of QatB are colored based on plDDT scores.

**b,** Comparison of structure of the QatBCD complex predicted by Alphafold3 and solved QatBC complex. The N-terminus of QatB is represented by a blue ellipse in two structures.

**c,** Structural alignment of the top five predicted structures of QatBCD complex. QatB and QatC are colored as in a. Two views are shown.

**d,** Native-PAGE showed the binding among QatB$^{\Delta 2\text{-}46}$, QatC and QatD. The results of the SDS-PAGE for the same sample are displayed on the right.

Extended Data Table 1

| Statistics | Apo | ATP |
|---|---|---|
| **Data collection** | | |
| Space group | P 21 | I 2 2 2 |
| Cell dimensions | | |
| *a*, *b*, *c* (Å) | 58.388 116.179 120.324 | 116.53 120.43 121.54 |
| (°) | 90 103.01 90 | 90 90 90 |
| Resolution (Å) | 58.62 - 2.09 (2.11 - 2.09)[a,b] | 38.12 - 2.66 (2.77 - 2.66)[a,b] |
| $R_{sym}$ or $R_{merge}$ (%) | 21.7 (122.0) | 15.41 (175.6) |
| $I/\sigma(I)$ | 5.9 (1.7) | 11.68 (1.38) |
| Completeness (%) | 100 (100) | 99.59 (99.30) |
| Redundancy | 6.5 (5.7) | 13.5 (12.9) |
| **Refinement** | | |
| Resolution (Å) | 58.62- 2.09 (2.11 - 2.09) | 38.12 - 2.66 (2.77 - 2.66) |
| Unique reflection | 92252 (2892) | 24800 (2687) |
| $R_{work}$ / $R_{free}$[c] | 0.219 /0.247 | 0.203/0.243 |
| No. atoms | 11752 | 5572 |
| Protein | 1409 | 5523 |
| Ligand/ion | 2 | 34 |
| Water | 953 | 15 |
| *B* factors | 28.63 | 75.38 |
| Protein | 28.07 | 75.40 |
| Ligand/ion | 27.86 | 77.51 |
| Water | 34.95 | 61.98 |
| R.m.s. deviations | | |
| Bond lengths (Å) | 0.002 | 0.002 |
| Bond angles (°) | 0.49 | 0.48 |
| Ramachandran plot (%) | | |
| Favored | 98.49 | 98.47 |
| Allowed | 1.51 | 1.53 |
| Outliers | 0 | 0 |

[a]For each structure one crystal was used.
[b]Values in parentheses are for highest resolution shell.
[c]Rfree was calculated with 5% of the reflections selected.
